# Accurate single-molecule spot detection for image-based spatial transcriptomics with weakly supervised deep learning

**DOI:** 10.1101/2023.09.03.556122

**Authors:** Emily Laubscher, Xuefei (Julie) Wang, Nitzan Razin, Tom Dougherty, Rosalind J. Xu, Lincoln Ombelets, Edward Pao, William Graf, Jeffrey R. Moffitt, Yisong Yue, David Van Valen

## Abstract

Image-based spatial transcriptomics methods enable transcriptome-scale gene expression measurements with spatial information but require complex, manually-tuned analysis pipelines. We present Polaris, an analysis pipeline for image-based spatial transcriptomics that combines deep learning models for cell segmentation and spot detection with a probabilistic gene decoder to quantify single-cell gene expression accurately. Polaris offers a unifying, turnkey solution for analyzing spatial transcriptomics data from MERFSIH, seqFISH, or ISS experiments. Polaris is available through the DeepCell software library (https://github.com/vanvalenlab/deepcell-spots) and https://www.deepcell.org.

## 1 Introduction

Advances in spatial transcriptomics have enabled system-level gene expression measurement while preserving spatial information, enabling new studies into the connections between gene expression, tissue organization, and disease states ^1,2^. Spatial transcriptomics methods fall broadly into two categories. Sequencing-based methods leverage arrays of spatially barcoded RNA capture beads to integrate spatial information and transcriptomes ^3–6^. Image-based methods, including multiplexed RNA fluorescent in situ hybridization (FISH) and in situ RNA sequencing (ISS), perform sequential rounds of fluorescent staining to label transcripts to measure the expression of thousands of genes in the same sample ^7–11^. Because these methods rely on imaging, the data that they generate naturally contain the sample’s spatial organization. While image-based spatial transcriptomics enables measurements with high transcript recall and subcellular resolution ^1,2^, rendering the raw imaging data interpretable remains challenging. Specifically, the computer vision pipelines for image-based spatial transcriptomics must reliably perform cell segmentation, spot detection, and gene assignment across diverse imaging data. Prior methods that sought an integrated solution to this problem rely on manually-tuned algorithms to optimize performance for a particular sample or spatial transcriptomics assay ^12,13^. Thus, there remains a need for an integrated, open-source pipeline that can perform these steps reliably across the diverse images generated by spatial transcriptomics assays with minimal human intervention.

Deep learning methods are a natural fit for this problem. Prior work by ourselves and others has shown that deep learning methods can accurately perform cell segmentation with minimal user intervention ^14–17^, providing a key computational primitive for cellular image analysis. Here, we focus on the problem of spot detection for image-based spatial transcriptomics data. Existing spot detection methods fall into two categories: “classical” and “supervised” ^18^. Classical methods are widely used but require manual parameter fine-tuning to optimize performance ^19,20^. The optimal parameter values are often different within regions of the same image, making implementation of classical methods time-intensive and fundamentally limiting their scalability. Supervised methods ^21–23^, which often rely on deep learning methodologies, learn how to detect spots from labeled training data. These methods eliminate the need for manual parameter tuning to optimize spot detection performance. However, the requirement for labeled training data presents a major challenge, as experimentally-generated data contain too many spots for manual annotation to be feasible. Training data derived from classical algorithms are limited by the characteristics of those algorithms, imposing a ceiling on model performance. Further, simulated training data lack the artifacts present in experimentally-generated data which can limit the model’s performance on real data.

In this work, we combine deep learning a with weakly supervised training data construction scheme to create a universal spot detector for image-based spatial transcriptomics data. We demonstrate the performance of our spot detection model on simulated and experimentally-generated images. Given that training deep learning models with weak supervision can yield a computational primitive for spot detection, we then constructed Polaris, an integrated deep learning pipeline for image-based spatial transcriptomics. Constructed in this fashion, Polaris offers a turnkey analysis solution for data from various image-based spatial transcriptomics methods while removing the need for manual parameter tuning or extensive user expertise.

## 2 Training spot detection models with weak supervision

Here, we describe two key aspects of Polaris’ spot detection model - constructing consensus training data with annotations from multiple classical spot detection algorithms, and deep learning model design and training.

### Constructing consensus training data

Accurate training data is an essential component of every deep learning method. In this work, we have sought to create training data for spot detection models by finding consensus among several commonly used classical spot detection algorithms (Fig. 1a). In our approach, we first create annotations for representative fluorescent spot images by manually fine-tuning a collection of classical algorithms on each image. We refer to these algorithms as “annotators”. This process generates conflicting sets of coordinate spot locations, as each annotator detects or misses different sets of spots. To determine which annotators detected each spot in an image, the detections from all annotators are clustered based on their proximity. Inspired by prior work on programmatic labeling ^24^, we then de-noise conflicting spot annotations with a generative model. The generative model characterizes annotators with two parameters: (1) true positive rate (TPR), which is an annotator’s probability of detecting a ground-truth true spot, and (2) false positive rate (FPR), which is an annotator’s probability of detecting a ground-truth false spot. The model characterizes clustered spots by their probability of corresponding to a ground-truth true spot (*p*(TP)). The generative model is given an initial guess for the TPR and FPR of each classical algorithm and a matrix of annotation data, which we name the “detection information matrix”. This matrix of annotation data, 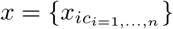 consists of binary variables, *x*_*ic*_, which are equal to 1 if annotator *i* detected cluster *c*, and 0 if not. The model is then fit with expectation maximization (EM) ^25^ by iteratively calculating the *p*(TP) of each cluster and estimating the TPRs and FPRs of each annotator until convergence (Fig. 1d). The remainder of this section provides more detail on the mathematical execution of these steps.

**Figure 1:**
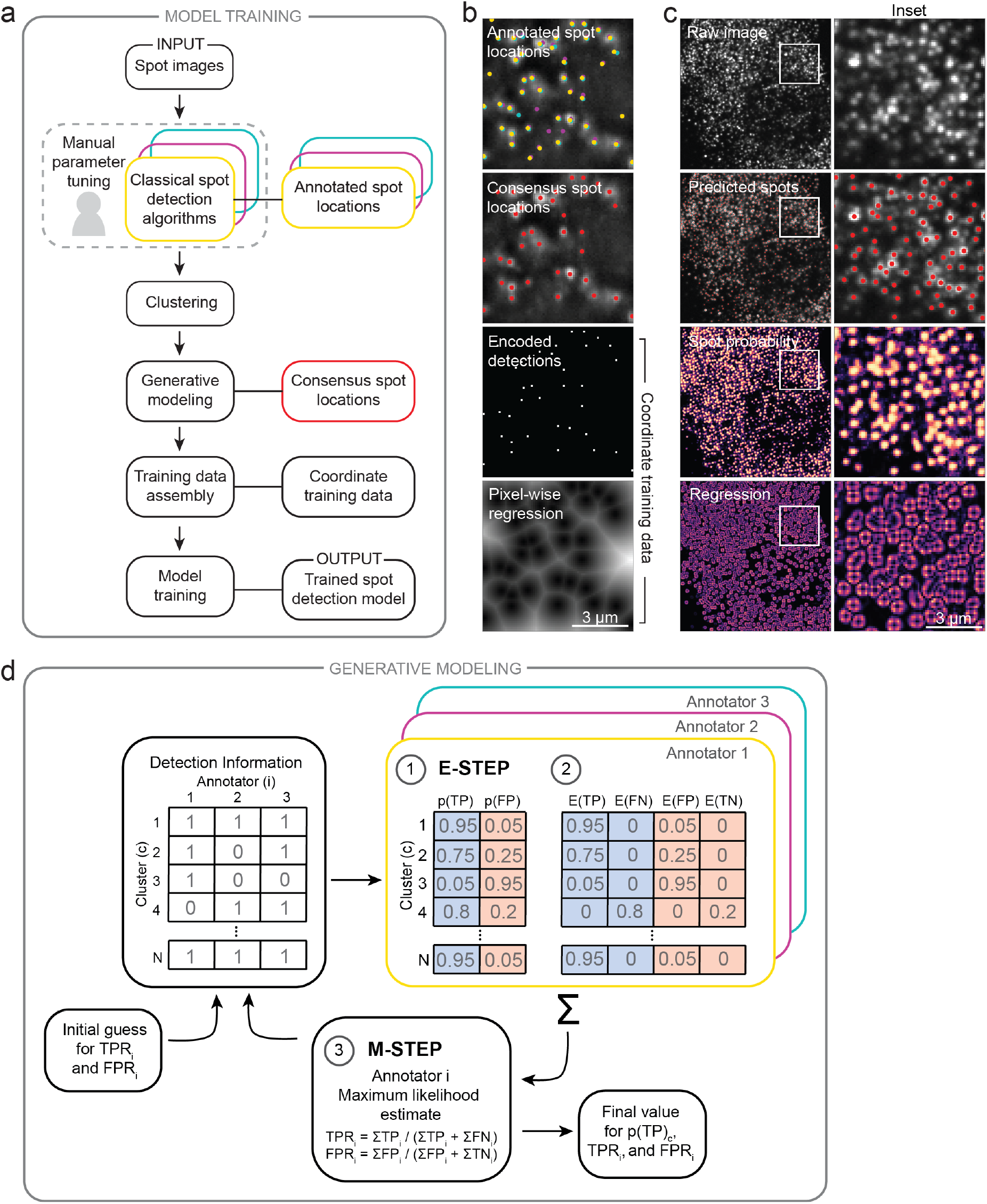
A weakly supervised deep learning framework for accurate fluorescent spot detection of spatial transcriptomics imaging data. (a) Training data generation for spot detection. Spot labels were generated by finding consensus among a panel of commonly used classical spot detection algorithms through generative modeling. These consensus labels were then used to train Polaris’ spot detection model. Sequential steps are linked with an arrow; associated methods and data types are linked with a solid line. (b) Demonstration of the training data generation for an example spot image. Spot locations are converted into encoded detections and distance maps which guide the classification and regression tasks performed during model training. Spot colors correspond to the annotation colors in (a). (c) Output of Polaris’ spot detection model for an example seqFISH image. Regression values above a default threshold are set to zero. The regression images in (b-c) are the sum of the squared pixel-wise regression in the x- and y-directions. (d) Schematic diagram of the EM method to fit the generative model for consensus spot annotation creation.

Our generative model assumes that each annotator *i* produces Bernoulli-distributed annotations. We define *z*_*c*_ to be a binary variable indicating whether cluster *c* is a true spot or not (1 if true, 0 if not). *p*(*z*_*c*_) - conditioned on the data and annotator characteristics - corresponds to *p*(TP), which we wish to compute. Let us also define *θ*_*i*_(*z*_*c*_) to be a variable that represents an annotator *i*’s probability of detecting a cluster, conditioned on whether the cluster is a true spot or not. This notation is a more compact way of representing the annotator characteristics, as *θ*_*i*_(1) = TPR_*i*_ and *θ*_*i*_(0) = FPR_*i*_. For every cluster *c* and annotator *i*, the distribution of *x*_*ic*_ given the cluster assignment *z*_*c*_ and annotator characteristics *θ*_*i*_(*z*_*c*_) is a Bernoulli distribution:

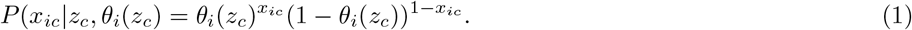

We assume the variables *x*_*ic*_ are independent; the probability to observe the data *x* = *{x*_*ic*_*}*, given *{θ*_*i*_*}* and *{z*_*c*_*}* is then

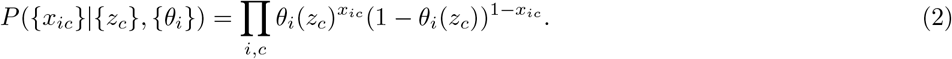

To offer a concrete example of this formula in action, consider the following situations for a hypothetical set of three annotators. For a “true detection” (e.g., *z*_*c*_ = 1), the probability that all three annotators detect the spot (e.g., *x*_*ic*_ = 1 for all *i*) given by the above formula reduces to

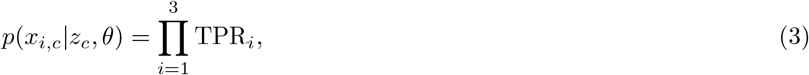

which is simply the product of the TPRs of each annotator. Alternatively, the probability that the first two annotators detect a ground-truth true spot while the third annotator (incorrectly) does not is given by:

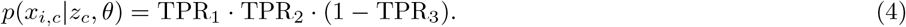

We utilize formula 2 with the EM algorithm to infer the annotator and cluster characteristics. The EM algorithm consists of two computation steps: an expectation step and a maximization step. To perform the expectation step, we define the probability of a cluster corresponding to a true or false detection with Bayes’ theorem:

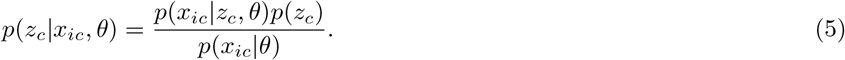

The term *p*(*z*_*c*_) is the prior probability of a cluster corresponding to a true or false detection; we use the least informative value for the prior by setting *p*(*z*_*c*_) = 1*/*2, indicating an equal probability that a spot is a true or false detection. The term *p*(*x*_*ic*_|*θ*) can be expressed as:

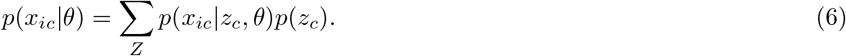

Therefore, the probability *p*(*z*_*c*_|*x*_*ic*_, *θ*) can be expressed as the likelihood of each possible label (e.g., true or false) normalized by the sum of the likelihood of both labels:

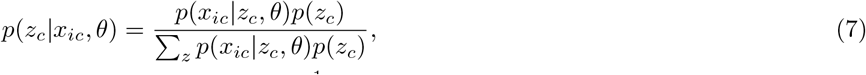

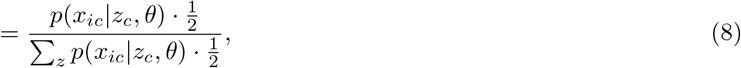

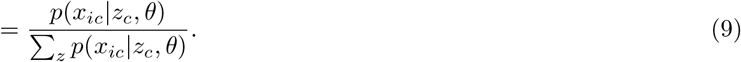

Using this method to calculate *p*(*z*_*c*_|*x*_*ic*_, *θ*), we can then calculate 𝔼 (TP), 𝔼 (FN), 𝔼 (FP), and 𝔼 (TN) for each annotator.

1. Two scenarios can arise when calculating these values.
2. If an annotator detects a spot in a particular cluster, i.e. *x*_*ic*_ = 1, 𝔼 (TP) for that annotator is equal to *p*(TP) = *p*(*z*_*c*_|*x*_*ic*_, *θ*) for that cluster, and 𝔼 (FP) for that annotator is equal to *p*(FP) for that cluster. 𝔼 (TN) and 𝔼 (FN) are set to zero.

If an annotator does not detect a spot in a particular cluster, i.e. *x*_*ic*_ = 0, 𝔼 (TN) for that annotator is equal to *p*(FP) = 1 *− p*(TP) for that cluster, and 𝔼 (FN) for that annotator is equal to *p*(TP) for that cluster. 𝔼 (TP) and 𝔼 (FP) are set to zero.

To perform the maximization step, we sum 𝔼 (TP), 𝔼 (FN), 𝔼 (FP), and 𝔼 (TN) across all clusters to calculate an updated maximum likelihood estimate for *TPR*_*i*_ and *FPR*_*i*_ for method *i* with equations of the following form:

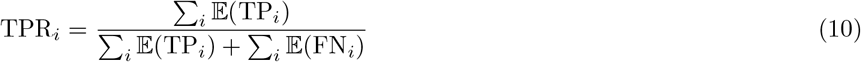

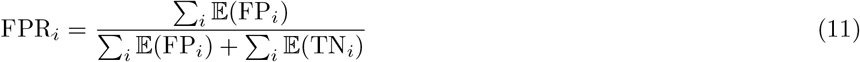

The expectation and maximization steps are performed iteratively until the values for *p*(TP)_*c*_, TPR_*i*_, and FPR_*i*_ converge. The consensus spot locations are taken at the centroid of the clusters with a *p*(TP)_*c*_ value that exceeds a defined probability threshold. We demonstrated that this method yields accurate estimation of TPR and FPR and that the resulting spot labels approach 100% correct as data set size and numbers of annotators increase (Supplementary Fig. S1).

To construct a training dataset for Polaris’ spot detection model, we applied this consensus training data construction method to images from various spatial transcriptomics assays. We assembled a set of representative images from sequential fluorescence in situ hybridization (seqFISH), multiplexed error-robust FISH (MERFISH) ^26,27^, and SunTag-labeled imaging data ^28^. We use these consensus annotations to train a deep learning model for spot detection.

### Model design and training

To train our spot detection model, we frame the problem as a classification and regression task (Fig. 1b). For each pixel we seek to predict whether that pixel contains a spot and compute the distance from the pixel to the nearest spot centroid. To train our model with our consensus spot labels, the coordinate spot locations are converted into two image types: (1) an image containing one-hot encoded spot locations, and (2) regression images encoding the sub-pixel distance to the nearest spot in the x- and y-directions (Fig. 1b). The deep learning model returns two output images from a given input image: (1) the pixel-wise probability of a spot, and (2) x- and y-regression images (Fig. 1c). Our model is trained with a weighted cross entropy and a custom mean squared error loss (e.g., computed only in a neighborhood around each spot) for the two output images (see Supplementary Note 2). For our model architecture, we utilize FeatureNets, a family of models that are parameterized by their receptive field ^14^. We perform hyperparameter optimization experiments to find the optimal receptive field size of 13 (Supplementary Fig. S5). To return the location of the spots in an image, we use maximum intensity filtering to detect the local maxima in the spot probability image and use the regression images to update the coordinates of each spot to achieve sub-pixel resolution.

## 3 Results

We demonstrate Polaris’ spot detection capabilities on held-out experimentally-generated images. Visual inspection showed that our model generalized to out-of-distribution, spot-like data generated by various spatial transcriptomics assays, such as ISS ^7^ and splitFISH images ^11^ (Supplementary Fig. S2). Additionally, we used held-out images to quantify the agreement between Polaris and the classical methods used to create our consensus training data. Agreement between sets of detected spots was determined with a mutual nearest neighbors matching method. (Supplementary Fig. S3). We observed higher agreement between Polaris and the classical methods than exists among the classical methods themselves. This analysis demonstrates Polaris’ learning of consensus labels generalizes to images held-out from the training data set. (Supplementary Fig. S4)

The ambiguity of ground-truth annotations for experimental data presents challenges for quantitatively benchmarking spot detection methods. To evaluate the accuracy of our approach, we followed prior work by simulating spot images, which have unambiguous ground truth spot locations. ^29,30^ When accurate simulation of experimental data is possible, simulations remove the need for unambiguous ground-truth annotations for benchmarking. We note that our spot simulations add signal on top of autofluorescence images. Because we control the image generation, we can explore model performance as a function of image difficulty by tuning parameters such as the spot density and signal-to-noise ratio. Benchmarking on simulated data demonstrated that our method outperforms models trained with either simulated data or data labeled with a single classical algorithm. We found that this performance gap held across the tested range of spot intensity and density (Supplementary Fig. S6a-b). We concluded that the consensus annotations more accurately capture the ground truth locations of spots in training images than any single classical algorithm and that there is significant value to training with experimentally generated images. We also found that Polaris’ spot detection model outperforms other recently published spot detection methods when evaluated on these simulated data, demonstrating greater robustness to ranging spot density and signal-to-noise ratios (Supplementary Fig. S6c-d). The combination of benchmarking on simulated data, visual inspection, and analysis of inter-algorithm agreement led us to conclude that Polaris can accurately perform spot detection on a diverse array of challenging single-molecule images.

Polaris packages this model into an analysis pipeline implemented in Python for multiplexed spatial transcriptomics data. Polaris integrates multiple analysis steps to yield coordinate spot locations with assigned gene identities. (Fig. 2a) First, it utilizes classical computer vision methods for image alignment and performs spot detection on images across all staining rounds. Then, cell segmentation is performed with models from DeepCell software library ^16^. Polaris’ spot detection model predicts the pixel-wise spot probability for each imaging round. For multiplexed spatial transcriptomics images, Polaris considers a codebook of up to thousands of barcodes that define the rounds and colors of fluorescent staining for each gene. To assign gene identities for barcoded spatial transcriptomics images, we fit a graphical model of a mixture of relaxed Bernoulli distributions to the pixel-wise probability values with variational inference ^31,32^ (Fig. 2b-c). This model estimates the characteristic relaxed Bernoulli distributions of pixel values for “spots” and “background” in each imaging round, which may vary due to factors such as fluorophore identity, staining efficiency, or image normalization. These distributions are used in combination with the experimental codebook to estimate the probability of each barcode identity and ultimately assign spots to a gene identity or “background”.

**Figure 2:**
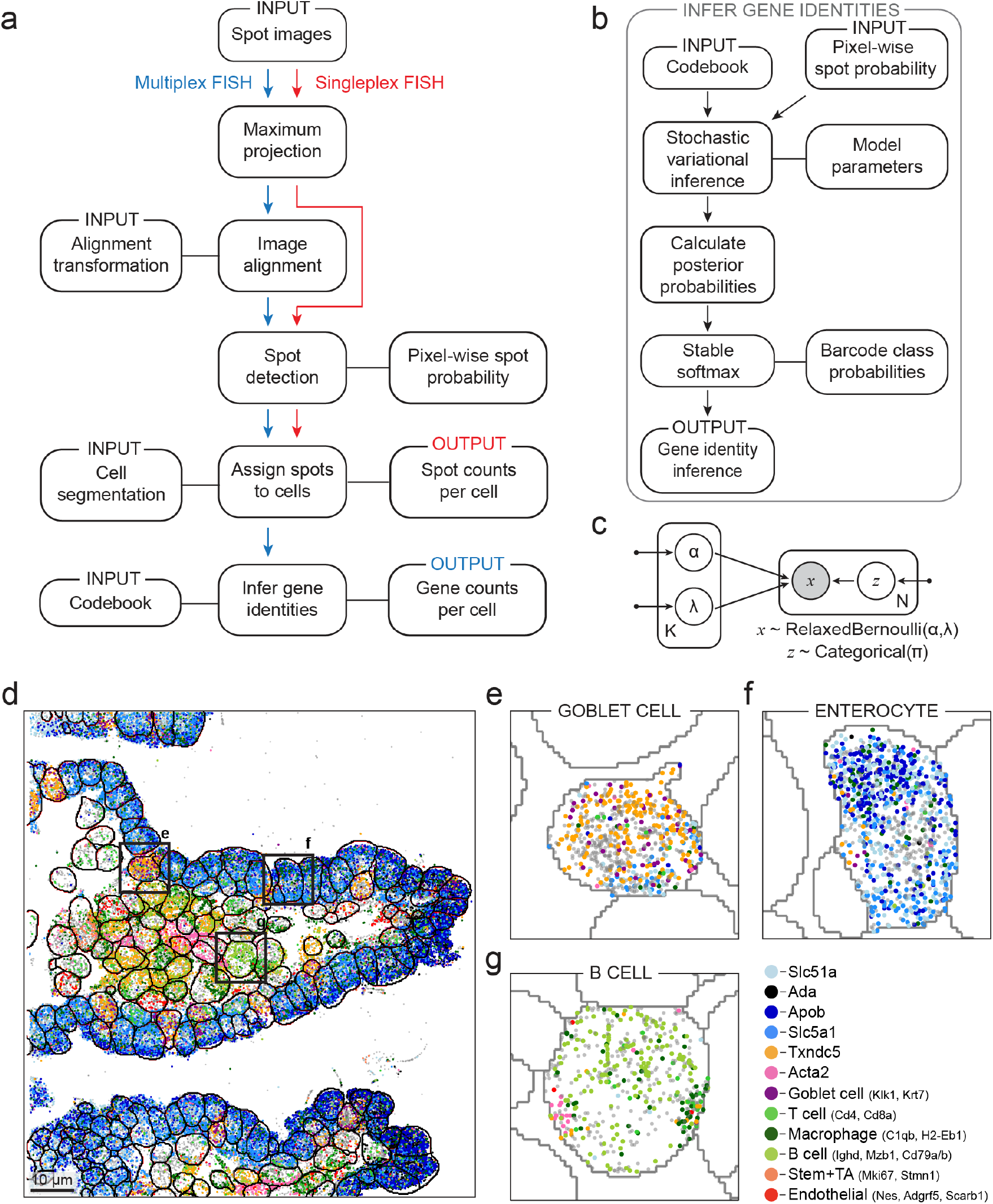
Polaris produces single-cell, spatial gene expression maps for multiplex spatial transcriptomics images. (a, b) Analysis steps for Polaris for singleplex (red) and multiplex (blue) spatial transcriptomics imaging data. Sequential steps are linked with an arrow, and associated methods and data types are linked with a solid line. Deep learning models perform spot detection and cell segmentation, while a probabilistic graphical model infers gene identities. (c) A probabilistic graphical model for inferring gene identities from spot detections. This model consists of a mixture of *K* relaxed Bernoulli distributions, parameterized by their probability, *α*, and their temperature, *λ*, for generating observed data, *x*, of size *N* spots. Shaded vertices represent observed random variables; empty vertices represent latent random variables; edges signify conditional dependency; rectangles (“plates”) represent independent replication; and small solid dots represent deterministic parameters. (d) Spatial organization of marker gene locations in a mouse ileum tissue sample. Each spot corresponds to a decoded transcript for a cell type marker gene. Whole-cell segmentation was performed with Mesmer ^16^. (e-g) Locations of decoded genes in an example Goblet cell, enterocyte, and B cell, respectively.

As with our benchmarking of spot detection methods, we used simulated data to benchmark the performance of our barcode assignment method quantitatively. Here, simulated data allowed us to explore our method’s dependency on spot dropout. This event can occur due to labeling failure, image quality, or failure in spot detection. Regardless of origin, the presence of dropout imposes a robustness constraint on the gene decoding methodology, as decoding schemes robust to dropout would better tolerate labeling and spot detection model failures. Our benchmarking of spot decoding with simulated data demonstrates that decoding with a generative model based on the relaxed Bernoulli distribution was more robust to dropout than other benchmarked methods (Supplementary Fig. S7).

We demonstrated Polaris’ performance on a variety of previously published data: a MERFISH experiment in a mouse ileum tissue sample ^27^ (Fig. 2d-f), a MERFISH experiment in a mouse kidney tissue sample ^33^ (Supplementary Fig. S8a), a seqFISH experiment in cultured macrophages (Supplementary Fig. S8b), and an ISS experiment of a pooled CRISPR library in HeLa cells ^34^ (Supplementary Fig. S10). We found that Polaris detected marker genes from expected cell types - even in areas with high cell density and heterogeneous cell morphologies in tissue samples. These results highlight the power of spatial transcriptomics methods to quantify gene expression while retaining multicellular and sub-cellular spatial organization.

For both tissue MERFISH datasets, we found that Polaris’ output gene expression counts have similar correlations with bulk sequencing data as the original analyses (r=0.796 and r=0.683; r=0.537 and r=0.565), and for both datasets, the two analysis outputs were highly correlated (r=0.936; r=0.911) (Supplementary Fig. S9a-d). For the cell culture seqFISH dataset, Polaris’ output has a similar correlation with bulk sequencing data as the output of the original analysis tool (r=0.809 and r=0.694) and the two outputs are highly correlated (r=0.910) (Supplementary Fig. S9e-f). For the ISS dataset, the barcode counts quantified with Polaris and the original analysis ^34^ are highly correlated (r=0.946) with Polaris consistently yielding higher counts (Supplementary Fig. S10). Spatial transcriptomics methods often encounter overdispersion in measuring gene expression counts, potentially limiting the efficacy of comparing these counts using a linear regression model. ^33,35^. Despite this limitation, these results demonstrate that Polaris can generalize across sample types, imaging platforms, and spatial transcriptomics assays without manual parameter tuning.

## 4 Discussion

We sought to create a key computational primitive for spot detection and an integrated, open-source pipeline for image-based spatial transcriptomics. Our weakly supervised deep learning model for spot detection provides a universal spot detection method for image-based spatial transcriptomics data. Our training data generation methodology effectively tackles a fundamental data engineering challenge in generating annotations for supervised spot detection methods, surpassing the performance achieved by using simulated data or a single classical method. Polaris packages this model and others into a unified pipeline that takes users from raw data to interpretable spatial gene expression maps with single-cell resolution. We believe Polaris will help standardize the computational aspect of image-based spatial transcriptomics, reduce the amount of time required to go from raw data to insights, and facilitate scaling analyses to larger datasets. Polaris’ outputs are compatible with downstream bioinformatics tools, such as squidpy ^36^ and Seurat ^37^. Polaris is available for academic use through the DeepCell software library https://github.com/vanvalenlab/deepcell-spots and as a Python package distributed on PyPI https://pypi.org/project/DeepCell-Spots/. A singleplex deployment of the pipeline is available through the DeepCell web portal https://deepcell.org.

## Acknowledgments

We thank Lior Pachter, Barbara Englehardt, Sami Farhi, Ross Barnowski, and the other members of the Van Valen lab for useful feedback and interesting discussions. We thank Nico Pierson and Jonathan White for contributing data and providing early annotations. The HeLa cell line was used in this research. Henrietta Lacks, and the HeLa cell line established from her tumor cells without her knowledge or consent in 1951, have made significant contributions to scientific progress and advances in human health. We are grateful to Henrietta Lacks, now deceased, and her surviving family members for their contributions to biomedical research. This work was supported by awards from the Shurl and Kay Curci Foundation (to DVV), the Rita Allen Foundation (to DVV), the Susan E Riley Foundation (to DVV), the Pew-Stewart Cancer Scholars program (to DVV), the Gordon and Betty Moore Foundation (to DVV), the Schmidt Academy for Software Engineering (to TD), the Michael J Fox Foundation through the Aligning Science Across Parkinsons consortium (to DVV), the Heritage Medical Research Institute (to DVV), the NIH New Innovator program (DP2-GM149556) (to DVV), and an HHMI Freeman Hrabowski Scholar award (to DVV).

## Author Contributions

EL, NR, and DVV conceived the project. EL, NR, and DVV conceived the weakly supervised deep learning method for spot detection. EL and EP created the seqFISH training data for the spot detection model. LO contributed to the seqFISH protocol used to create training data. EL developed software for training data annotation. EL curated and annotated the training data for the spot detection model. NR and EL developed the spot detection model training software. EL trained the models. EL and NR developed the metrics software for the spot detection model. XW, EL, YY, and DVV conceived the combinatorial barcode assignment method. EL and XW developed the barcode assignment software, with input from YY and DVV. EL developed the multiplex image analysis pipeline. EL and XW benchmarked the multiplex image analysis pipeline. EL and TD developed the cloud deployment. RJX and JRM collected and analyzed MERFISH data. EL and EP created the macrophage seqFISH dataset. WG and DVV oversaw software engineering for Polaris. YY and DVV oversaw the algorithm development for the project. EL and DVV wrote the manuscript, with input from all authors. DVV supervised the project.

## Code availability

Code used for cell segmentation and model development is available at https://github.com/vanvalenlab/deepcell-tf. The Python package containing the spot detection and gene assignment code is available on Github https://github.com/vanvalenlab/deepcell-spots or PyPI https://pypi.org/project/DeepCell-Spots/. An example Jupyter notebook for Polaris is available at https://github.com/vanvalenlab/deepcell-spots/blob/master/notebooks/applications/Polaris-application.ipynb. The code used for model deployment is available at https://github.com/vanvalenlab/kiosk-console. Finally, code for reproducing all models and figures included in the paper is available at https://github.com/vanvalenlab/Polaris-2023_Laubscher_et_al.

## Data availability

The data and annotations used to train the spot detection model are available for academic use at https://deepcell.readthedocs.io/en/master/data-gallery.

## 5 Methods

### 5.1 Generation of sequential fluorescent in situ hybridization (seqFISH) images for spot training data

#### 5.1.1 Probe design

mRNA transcripts were targeted with single-stranded DNA probes, as previously described ^10^. Primary probes were designed to target a panel of 10 genes with OligoMiner using balanced coverage settings ^38^. The primary probes were designed to have secondary probe-binding sites flanking both ends of the sequence that binds to the mRNA transcript. The secondary probes were 15 bases long and consisted of nucleotide combinations that were optimized to have 40%-60% GC content and minimal genomic off-target binding.

#### 5.1.2 Probe construction

Single-stranded DNA primary probes were obtained from Integrated DNA Technologies (IDT) as an oPools Oligo Pool. Oligos were received lyophilized and dissolved in ultrapure water at a stock concentration of 1 *µ*M per probe. Single-stranded DNA secondary probes were also obtained from IDT and were 5’-functionalized with Alexa Fluor 647, Alexa Fluor 546, or Alexa Fluor 488. Secondary probes were received lyophilized and dissolved in ultrapure water at a concentration of 100 nM.

#### 5.1.3 Cell culture

HeLa (CCL-2) cells were received from the American Type Culture Collection. The cells were cultured in Eagle’s minimum essential medium (Cytiva #SH30024LS) supplemented with 2 mM L-glutamine (Gibco), 100 U/mL penicillin, 100 *µ*g/mL streptomycin (Gibco or Caisson), and 10% fetal bovine serum (Omega Scientific or Thermo Fisher). Cells were incubated at 37^*°*^C in a humidified 5% CO_2_ atmosphere and were passaged when they reached 70%-80% confluence.

#### 5.1.4 Buffer preparation

The primary probe hybridization buffer consisted of 133 mg/mL high-molecular-weight dextran sulfate (Calbiochem #3710-50GM), 2X saline-sodium citrate (SSC) (IBI Scientific #IB72010), and 66% formamide (Bio Basic #FB0211-500) in ultrapure water. The 55% wash buffer was comprised of 2X SSC, 55% formamide, and 1% Triton-X (Sigma-Aldrich #10789704001) in ultrapure water. The secondary probe hybridization buffer consisted of 2X SSC, 16% ethylene carbonate (Sigma-Aldrich #E26258-500G), and 167 mg/mL of high-molecular-weight dextran sulfate in ultrapure water. The ethylene carbonate was first melted at 50^*°*^C for 30-60 minutes. The 10% wash buffer was comprised of 2X SSC, 10% formamide, and 1% Triton-X in ultrapure water. The imaging buffer base consisted of 0.072M Tris HCl (pH 8) (RPI #T60050-1000), 0.43 M NaCl (Fisher #MK-7581-500), and 3 mM Trolox (Sigma-Aldrich #238813) in ultrapure water. The anti-bleaching buffer was comprised of 70% imaging buffer base, 2X SSC, 1% catalase (10X dilution of stock) (Sigma #C3155), 0.005 mg/mL glucose oxidase (Sigma-Aldrich #G2133-10KU), and 0.08% D-glucose (Sigma #G7528) in ultrapure water.

#### 5.1.5 seqFISH sample preparation

The cells were seeded in a fibronectin-functionalized (Fisher Scientific #33010018) glass-bottom 96-well plate (Cellvis P96-1.5H-N) at 80%-90% confluence. Cells were rinsed with warm 1X phosphate-buffered saline (PBS) (Gibco), fixed with fresh 4% formaldehyde (Thermo #28908) in 1X PBS for 10 minutes at room temperature, and then permeabilized with 70% ethanol overnight at −20^*°*^C. Prior to probe hybridization, the cells were rinsed with 2X SSC. The primary probes were diluted to 10 nM in the primary probe hybridization buffer and 100 *µ*L of this solution was added to each well. Cells were incubated with the primary probe solution for 24 hours at 37^*°*^C. The primary probes were rinsed out twice with 55% wash buffer. Cells were incubated in 55% wash buffer for 30 minutes in the dark at room temperature and then rinsed three times with 2X SSC buffer. The secondary probes were diluted to 50 nM in the secondary probe hybridization buffer and 100 *µ*L was added to each of the wells. The probes were then incubated for 15 minutes in the dark at room temperature. The secondary probes were then washed twice in 10% wash buffer. The cells were then incubated in the 10% wash buffer for 5 minutes in the dark at room temperature. Finally, the cells were washed once with 2X SSC buffer and once with imaging buffer. Before imaging, the buffer was changed to anti-bleaching buffer (100 *µ*L).

Images of cell autofluorescence were acquired with non-specific secondary probe staining to generate simulated spot images. These samples were prepared with the same seqFISH method described above, without the addition of primary probes.

#### 5.1.6 Imaging conditions

The seqFISH samples were imaged with a Nikon Ti2 fluorescence microscope controlled by Nikon Elements. Images were acquired with a Nikon SOLA SE II light source, a 100X oil objective, and a Photometrics Prime 95B CMOS camera.

### 5.2. Creation of spot training data

#### 5.2.1 Image annotation

Our training dataset consisted of 1000 128×128 pixel images: 400 images generated as described above by performing seqFISH on cell culture samples, 400 previously published images generated with multiplexed error-robust FISH on tissue samples ^26,27^, and 200 previously published images generated with SunTag labeling of nascent proteins in cell culture samples ^28^. All data were scaled so that the pixels had the same physical dimension of 110 nm prior to training. These images contained up to approximately 200 spots per image and were min-max normalized to a range of [0,1] prior to annotation. We annotated each image with five classical spot detection algorithms: maximum intensity filtering (`skimage.feature.peak local max`), difference of Gaussians (`skimage.feature.blob dog`), Laplacian of Gaussian (`skimage.feature.blob log`), the Crocker-Grier centroid-finding algorithm (`trackpy.locate`), and Airlocalize. To accelerate image labeling with all methods except Airlocalize, we created a Tkinter graphical user interface to tune the algorithm parameters (intensity threshold, minimum distance between spots, etc.) on a per-image basis. To create annotations with Airlocalize, we utilized its previously published Matlab GUI to fine tune the algorithm parameters on a per-image basis. ^39^

For the spot detection algorithms that return spot locations with pixel-level resolution (`peak local max`, `blob dog`, and `blob log` in `skimage.feature`), the subpixel localization was determined by fitting a 2D isotropic Gaussian to the spot intensity ^40^. A 10×10 pixel portion of the image surrounding the detected spot was cropped out and used for the Gaussian fitting. A nonlinear least squares regression was performed, with initial parameters of Gaussian mean at the pixel center, an amplitude of 1 (the image’s maximum value after min-max normalization) and a standard deviation of 0.5 pixels. The spot location was constrained to be in the middle 20% of the image, and the spot standard deviation was constrained to be between 0 and 3.

#### 5.2.2 Clustering of spot annotations

Spots detected by the four classical algorithms were clustered into groups by proximity. For clarity, we refer to these classical algorithms as “annotators”. Each group of detections was presumed to be derived from the same spot in the image. To perform this clustering, we first constructed a graph with each detected spot being a node. Two detections were connected by an edge if they were within 1.5 pixels of each other. We then considered the connected components of this graph to be clusters. We screened the clusters to ensure that they contained at most one detection from each algorithm. If a cluster contained more than one detection from the same algorithm, the detection closest to the cluster centroid was retained, and all other detections were separated into new clusters.

From this graph, we derived the “detection information matrix,” which identifies the annotators that contribute a detection to each cluster. This matrix has dimensions of *C · A*, where *C* is the number of connected components or clusters in the graph and *A* is the number of annotators. The matrix has a value of 1 or 0 when a particular annotator does or does not have a detection in a particular cluster, respectively.

#### 5.2.3 Creation of consensus annotations with expectation maximization

A generative model fit with the expectation-maximization (EM) algorithm ^25^ was used to estimate the probability that a cluster of detections corresponds to a true spot in the image. The detection information matrix, described above, was used as the input into the generative model along with an initial guess for the true positive rate (TPR) and false positive rate (FPR) of each algorithm and a prior probability of a spot being a true detection. The initial guesses for the TPR and FPR were 0.9 and 0.1, respectively. The prior probability of a spot being a true detection was defined as 0.5. Briefly, the EM algorithm consists of two steps - an expectation step and a maximization step. The expectation step yields an estimate for the probability that each detection cluster corresponds to a true detection. The maximization step yields an updated estimate for the TPR and FPR of each annotator. The expectation and maximization steps were performed iteratively 20 times, sufficient iterations for convergence to a local maximum of the likelihood. The resulting values were used as an estimate for the Bayesian probability of each cluster corresponding to a “true” spot. We provide further details of these steps in the main text. Clusters with a probability above 0.9 were used as spots in the training dataset, and the spot location was taken to be the centroid of the detection cluster.

### 5.3 Model benchmarking with simulated data

#### 5.3.1 Creation of simulated spot images

Simulated spot images were used to benchmark Polaris’ spot detection model. We created simulated images by adding Gaussian spots at random locations to cellular autofluorescence images. The location and number of spots in an image were sampled from uniform distributions. The intensity and width of the simulated Gaussian were sampled from normal distributions reflecting a spot distribution that is characteristic of experimental images.

#### 5.3.2 Creation of simulated detection information

Simulated detection information was used to benchmark the EM algorithm for generating consensus spot annotations. First, we simulated a set of spot identities, as either true detections or false detections. The ratio of ground-truth true and false spots was determined by a pre-defined prior probability of true spots. We then used the defined TPR and FPR values to simulate detections. The probability that true spots are detected by the simulated spot detection methods is defined by the simulated TPR, and the probability that false spots are detected is defined by the simulated FPR. As with detections from experimental images, the simulated detections are stored in a detection information matrix.

#### 5.3.3 Creation of simulated barcode pixel values

Simulated barcode pixel values were used to benchmark Polaris’ barcode assignment method’s robustness to dropout. We generated simulated barcode pixel values by sampling from distributions of spot and background pixel values from experimental images. The benchmarked methods include a graphical model of relaxed Bernoulli distributions, a graphical model of multivariate normal distributions, barcode matching by Hamming distance, and PoSTcode ^32^. The relaxed Bernoulli graphical model, the multivariate normal graphical model, and the distance matching method take input pixel values sampled from a distribution simulating spot probability values output by Polaris. Alternatively, PoSTcode takes input pixel values from a distribution simulating pixel values of the raw imaging data, consistent with its original methodology. Our method for simulating barcode values does not consider correlations in pixel values between images acquired in the same fluorescence channel or imaging round. PoSTcode relies on this nuanced characteristic of experimental data; thus, its performance on the simulated data used in this benchmarking analysis may have been negatively impacted.

#### 5.3.4 Mutual nearest neighbors point matching method

To quantitatively benchmark the performance of our deep learning models, we need a method for comparing sets of coordinate spot locations. Our method finds sets of mutual nearest neighbors to compare sets of ground-truth and predicted spot locations. To be considered a true detection, a detection must be within some threshold distance of a ground-truth detection. All spots outside this threshold distance are considered false detections (Supplementary Fig. S3a). If more than one detection is within the threshold distance of a ground-truth spot, the detection that is the closest to the ground-truth spot is considered a true detection, and all others are considered false detections (Supplementary Fig. S3b). If more than one detection is within the threshold distance for more than one ground-truth spot, edge cases may arise. For a detection to be considered a true detection, that detection and the corresponding ground-truth spot must be mutual nearest neighbors. Therefore, if a detection is within the threshold distance for two ground-truth spots, it is only paired with a ground-truth spot if they are each others’ mutual nearest neighbors. Otherwise, the detection is considered a false detection, even if the detection is within the threshold distance of a ground-truth spot (Supplementary Fig. S3c).

### 5.4. Spot detection deep learning model architecture

#### 5.4.1 Preparation of coordinate annotations for training data

The coordinate spot locations were converted into two different types of images before being used for deep learning model training. The first image type is a classification image array in which pixels corresponding to spots and background are one-hot encoded. The second image type is a regression image array, in which pixel values correspond to the distance to the nearest spot in the x- and y-direction.

#### 5.4.2 Image preprocessing

We performed preprocessing of images prior to model training. Pixel intensities were clipped at the 0.1 and 99.9th percentiles and then min-max normalized so that all pixel values were scaled between 0 and 1.

#### 5.4.3 Model architecture

Our deep learning model architecture is based on FeatureNets, a previously published backbone implemented in TensorFlow ^41^ where the receptive field is an explicit hyperparameter ^14^. We attach two prediction heads to this backbone: a classification head (to predict the probability a given pixel contains a spot) and a regression head (to predict the distance to the nearest spot with sub-pixel resolution). The receptive field of the network is an explicit hyperparameter that was set to a default value of 13 pixels.

All models were trained with stochastic gradient descent with Nesterov momentum. We used a learning rate of 0.01 and momentum of 0.9. We performed image augmentation during training to increase data diversity; augmentation operations included rotating (0^*°*^-180^*°*^), flipping, and scaling (0.8X-1.2X) input images. The labeled data were split into training and validation sets, with the training set consisting of 90% and the validation set consisting of 10% of the data. The test set for benchmarking model performance consisted of simulated spot images and held-out experimentally generated spot images.

#### 5.4.4 Image postprocessing

We processed the classification and regression predictions to produce a list of spots. The local maxima of the classification prediction output were determined using maximum intensity filtering, with a default intensity threshold of 0.95. For most datasets, we found that the spot detection results do not vary widely with changes to this threshold value. In the pixels determined to correspond to local maxima, the sub-pixel localization is determined by adding the value of the regression prediction in the x- and y-directions. These sub-pixel locations are returned as the output of the model.

### 5.5 Generation of multiplexed seqFISH dataset in cultured macrophages

#### 5.5.1 Cell culture

THP-1 (TIB-202) cells were received from the American Type Culture Collection. The cells were cultured in Roswell Park Memorial Institute (RPMI) 1640 Medium (Gibco) supplemented with 2 mM L-glutamine (Gibco), 100 U/mL penicillin, 100 *µ*g/mL streptomycin (Gibco or Caisson), and 10% fetal bovine serum (Omega Scientific or Thermo Fisher). To make the complete medium, 2-mercaptoethanol (BME) (Sigma-Aldrich #M6250) was added to a concentration of 0.05mM before every use. Cells were incubated at 37^*°*^C in a humidified 5% CO_2_ atmosphere and were passaged to maintain a concentration of 0.3-1 × 10^6^ cells/mL.

#### 5.5.2 seqFISH sample preparation and imaging

The THP-1 monocyte cells were seeded on a fibronectin-functionalized glass slide (Corning #2980-246) at 80%-90% confluence contained by a rubber gasket (Grace Bio-Labs #JTR8R-2.5). To differentiate the THP-1 monocytes into macrophages, the cells were incubated with 10ng/mL phorbol 12-myristate 13-acetate (PMA) (Sigma-Aldrich #P8139) in RPMI with BME for 24 hours. The media was then replaced with fresh RPMI with BME and incubated for an additional 24 hours. Differentiation was confirmed visually based on changes in adherence and morphology.

The macrophages were dosed with 1ug/mL LPS (Sigma Aldrich #L4524) in RPMI with BME for 3 hours. Cells were rinsed with warm 1X PBS, fixed with fresh 4% formaldehyde (Thermo #28908) in 1X PBS for 10 minutes at room temperature,and then permeabilized with 70% ethanol overnight at −20^*°*^C. The primary probe library (Spatial Genomics) was added to the sample in a flow chamber provided by Spatial Genomics and incubated overnight at 37^*°*^C. The sample was washed several times with primary wash buffer (Spatial Genomics). The nuclei of the sample were stained with staining solution (Spatial Genomics).

The macrophage sample was imaged with the Spatial Genomics Gene Positioning System (GenePS). Image tiling and secondary probe staining were performed programmatically by this instrument.

#### 5.5.3 Spatial Genomics image analysis

To generate a point of comparison for Polaris’ output, we analyzed the seqFISH dataset with the Spatial Genomics software. The spot detection for this analysis was performed via manual parameter tuning. For each imaging round and channel, the threshold intensity used to detect spots was defined visually. The validity of this threshold value was confirmed across several randomly selected fields of view. The DAPI channel was used as the input for their supervised nuclear segmentation method; nuclear masks were dilated to create whole-cell masks.

### 5.6 Multiplex FISH analysis pipeline

#### 5.6.1 Cell segmentation

For cell culture samples, cell segmentation was performed with nuclear and whole-cell segmentation applications from the Deepcell software library. For tissue samples, cell segmentation was performed with Mesmer ^16^. The source code for these models is available at https://github.com/vanvalenlab/deepcell-tf; a persistent deployment is available at https://deepcell.org.

#### 5.6.2 Gene identity assignment

Existing spot decoding methods for image-based spatial transcriptomics fall into two main categories: (1) pixel-wise decoding, which attempts to decode every pixel in the input image, and (2) spot-wise decoding, which attempts to detect spots before decoding them. Polaris’ spot decoding method uses elements of both methods. For spot decoding, Polaris’ pixel-wise spot probability output was used to determine which pixels to decode. The maximum intensity projection of the spot probability image was performed across all rounds and channels and the set of pixels to be decoded was determined by an intensity threshold with a default value of 0.01. For each thresholded pixel, the array of spot probability values at its coordinate location through the rounds and channels was used as the input for gene assignment.

Gene assignment was performed by fitting a generative model to the probability intensities at identified spot locations, a similar method to previously published work. ^32^ Our model consists of a mixture of 2 *· R · C* relaxed Bernoulli distributions, where *R* is the number of imaging rounds in the experiment and *C* is the number of fluorescent channels in each round. The model for pixel intensities was based on a mixture of relaxed Bernoulli distributions by default, but Polaris also offers two alternative distributions for modeling pixel intensities: Bernoulli and multivariate Gaussian. Therefore, the model consists of a “spot” distribution and a “background” distribution for each imaging round and channel. This model requires two inputs:

1. Spot probabilities at the pixel location across all imaging rounds and channels of the detected spots, as predicted by Polaris’ deep learning model
2. A codebook defining the imaging rounds and channels in which each gene in the sample is labeled, referred to as the gene’s barcode. An empty barcode is added to the input codebook, corresponding to a “background” - or false positive - assignment.

The distributions are fit to the pixel values of the detected spots with stochastic variational inference ^31^. The codebook is used to constrain the logit function of the relaxed Bernoulli distribution, and the temperature is learned. We assume independence across channels and imaging rounds, but the distribution parameters are shared across all genes. For each thresholded pixel, the probability of each barcode assignment is calculated, based on the probability that the pixel value was out of the “spot” or “background” distribution for each round and channel. These probability values are compared to the codebook to calculate the probability of each gene assignment for the pixel. The gene whose barcode has the highest probability is assigned to the pixel. If the highest probability assignment does not exceed a threshold probability value, set to 0.95 by default, the pixel is instead given an “unknown” assignment.

To find the coordinate locations of decoded genes, we create a mask with the pixels successfully decoded to a gene in the codebook and apply this mask to the maximum intensity-projected spot intensity image. We perform peak finding with maximum intensity filtering on the projected image to yield the coordinate location of each decoded gene.

#### 5.6.3 Gene assignment rescue methods

After prediction, “background” and “unknown” assignments can be rescued to be assigned gene identities through two methods. The first method, which we refer to as “error rescue”, compares the pixel values of the spot to each barcode in the codebook. If the Hamming distance of the pixel values to a barcode is less than or equal to 1, the assignment is updated. This method catches the rare cases in which the probabilistic decoder misses an assignment. The second method, which we refer to as “mixed rescue”, catches the spots that contain two mixed barcodes. This situation occurs when two RNA molecules are in close proximity in the sample and the signal from their barcode labels is mixed. Mixed barcodes often lead to a low probability assignment, so all spots below a threshold probability, set to 0.95 by default, are checked for this case. For this method, the barcode values for the original assignment are subtracted and the update pixel values are compared to each barcode in the codebook. If the Hamming distance of the pixel values to a barcode is less than or equal to 1, the assignment is updated.

#### 5.6.4 Background masking

Bright objects in the background of smFISH images can interfere with barcode assignment, so detected spots in these regions can be masked out. This step requires a background image without FISH staining to assess the initial fluorescence intensity in the sample. The background image is min-max normalized so that pixels are scaled between 0 and 1, and a mask is created with a threshold intensity, which defaults to 0.5. Any detected spots in the masked regions are excluded from downstream analysis. Polaris also allows a user to manually define a mask for image regions to exclude from analysis.

## 6 Supplementary Information

### 6.1 Supplementary Note 1: Evaluation of generative model performance for the creation of training data

**Figure S1:**
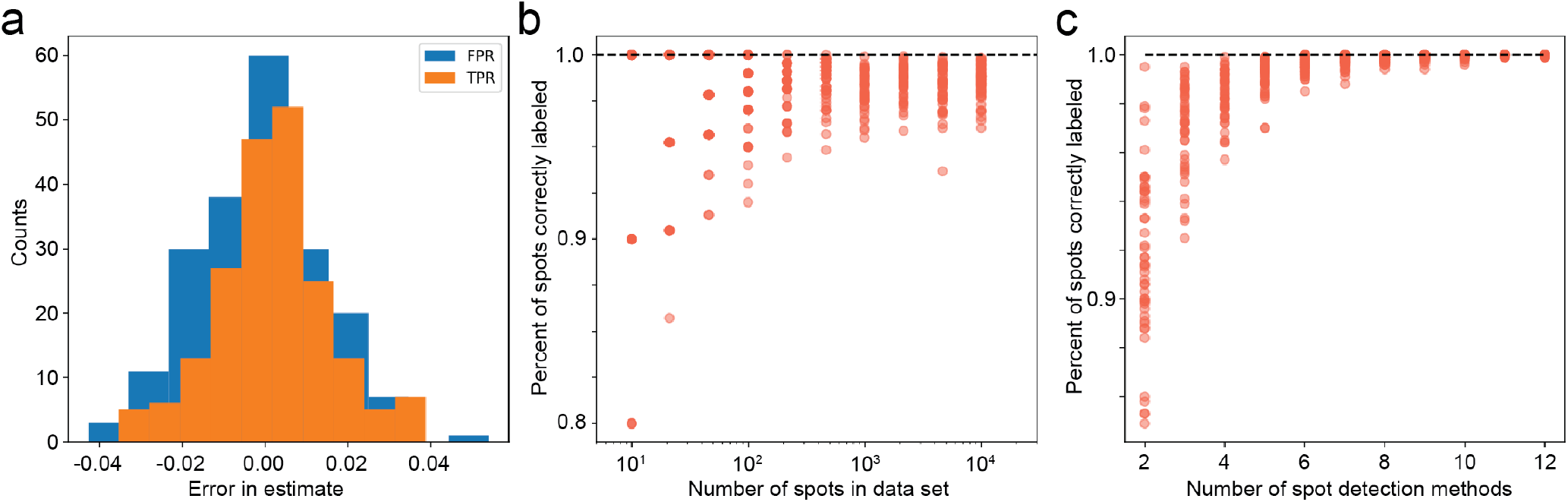
Benchmarking consensus annotation output of the generative model. (a) Error distribution for EM estimates of TPR and FPR values for 100 trials with three simulated classical methods. (b) Fraction of simulated detections correctly classified with increasing dataset size (number of spots in the dataset). (c) Fraction of simulated detections correctly classified as a true or false detection by EM for an increasing number of classical spot detection methods used in the EM method.

### 6.2 Supplementary Note 2: Custom loss function for Polaris’ deep learning model for spot detection

We trained the network using a custom loss function composed of a classification loss and a regression loss, which considers the outputs of both of the model’s prediction heads. The loss function has the following form:

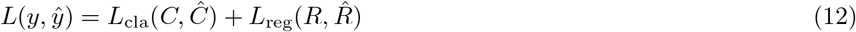

where *C* is the classification head output, *R* is the regression head output, and *y* = (*C, R*). The classification loss is the weighted cross-entropy with inverse class frequency-based weights. The regression loss is given by:

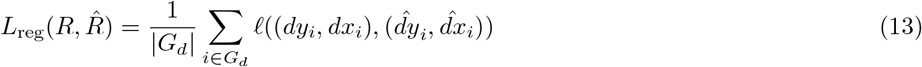

where *R*_*i*_ = (*dy*_*i*_, *dx*_*i*_) (*i* denotes a single pixel), *G*_*d*_ = *{*pixels *i* = (*i*_*y*_, *i*_*x*_)| a spot-containing pixel *j* exists with *L*_*∞*_(*i, j*) = max_*k*∈*x,y*_ |*i*_*k*_ *− j*_*k*_| ≤ *d}*, and *l* is the smooth *L*_1_ function. *dx*_*i*_ is the x-coordinate of the position difference between the nearest spot to pixel *i*, and pixel *i*’s center. Similarly, *dy*_*i*_ is the y-coordinate of this position difference. *d* is a configurable parameter that determines the threshold distance from the nearest spot under which the estimated nearest spot’s position for that pixel is taken into account in the loss function.

### 6.3 Supplementary Note 3: Generalization of Polaris’ spot detection model to a variety of spot images

**Figure S2:**
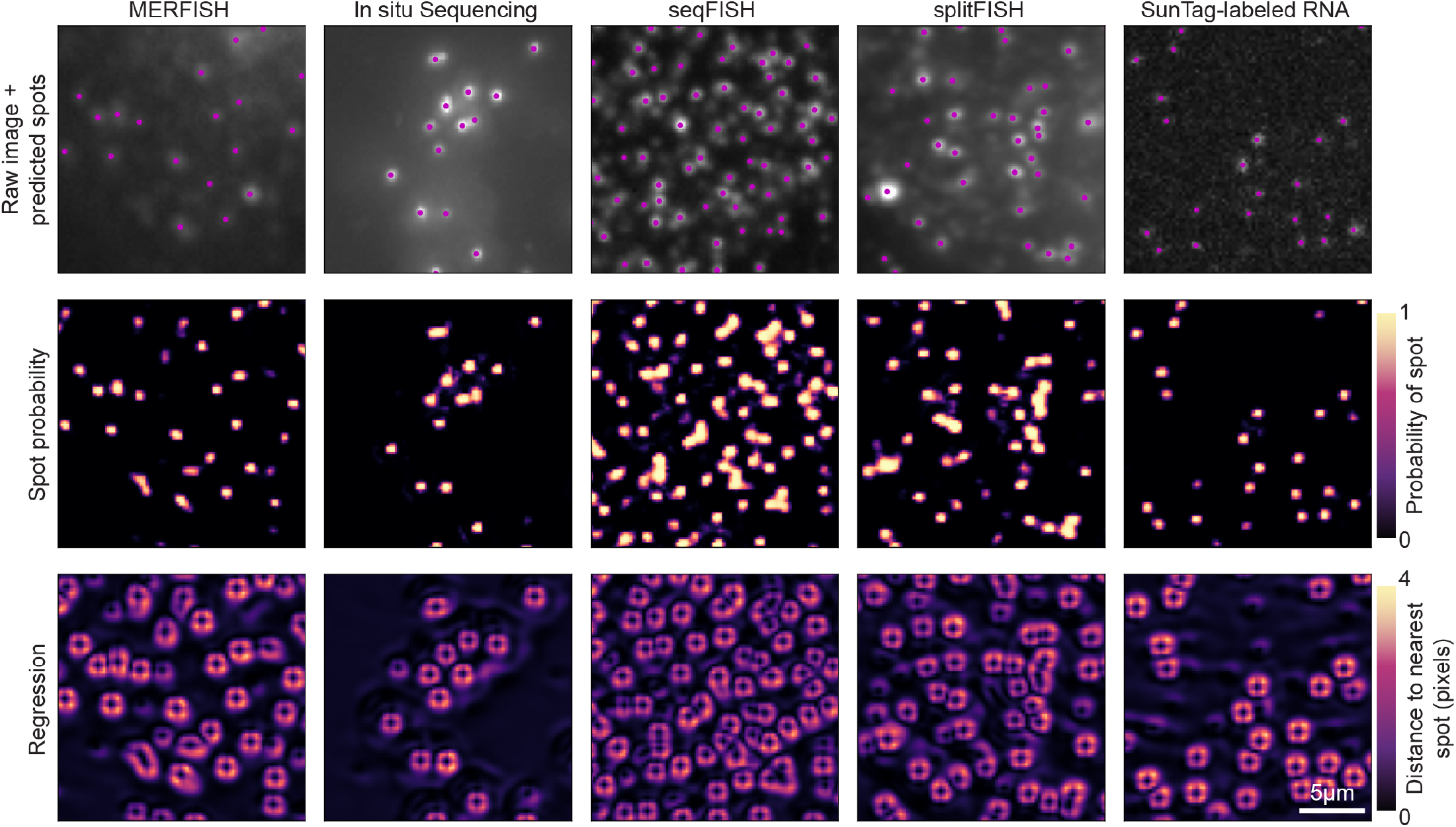
Polaris’ spot detection model generalizes to spot images generated with a variety of single-molecule assays. The spot probability prediction images encode the pixel-wise spot probability. The regression image is the sum of the square of the subpixel distances to the nearest spot in the x- and y-dimensions. Pixels beyond a threshold value are set to zero. These outputs are used together to generate a set of predicted spot locations with subpixel resolution, plotted over the raw image.

### 6.4 Supplementary Note 4: Mutual nearest-neighbor method for matching sets of spots

**Figure S3:**
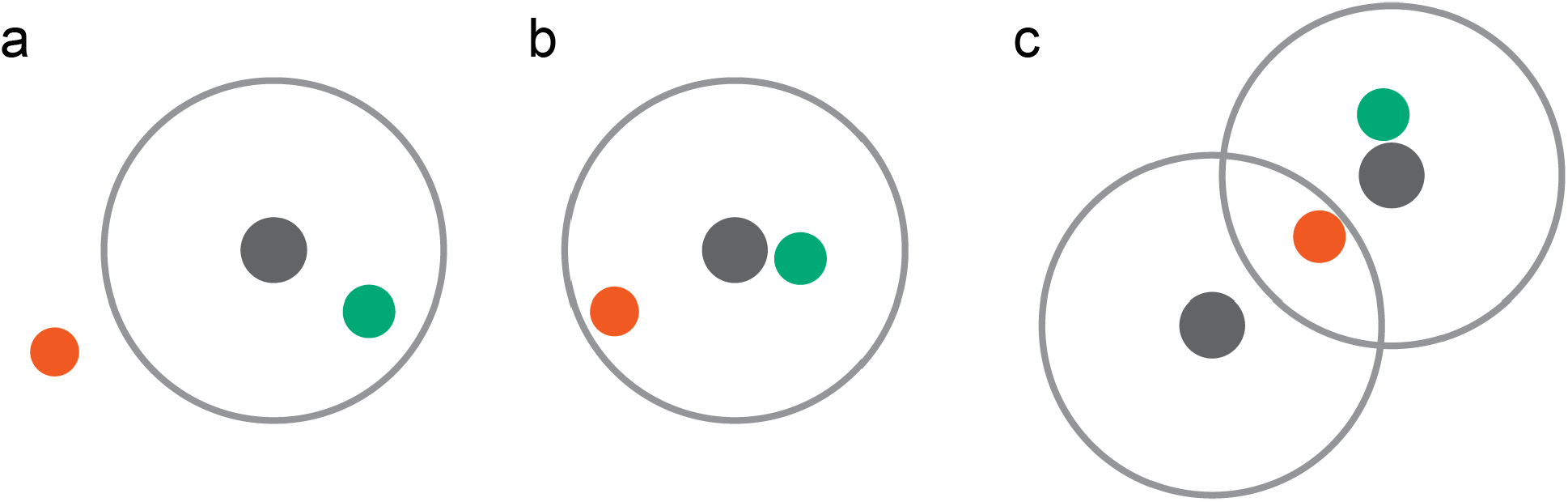
Example cases handled by a mutual nearest-neighbor matching algorithm. (a) Example with spots inside and outside the threshold distance to a ground-truth spot. Ground truth spots and their threshold distance are shown in grey. True positive detections are shown in green and false positive detections are shown in orange. (b) Example with two spots inside the threshold distance to a ground-truth spot. (c) Example with two spots within the threshold distance of two ground-truth spots.

### 6.5 Supplementary Note 5: Inter-algorithm agreement of spot detection results

**Figure S4:**
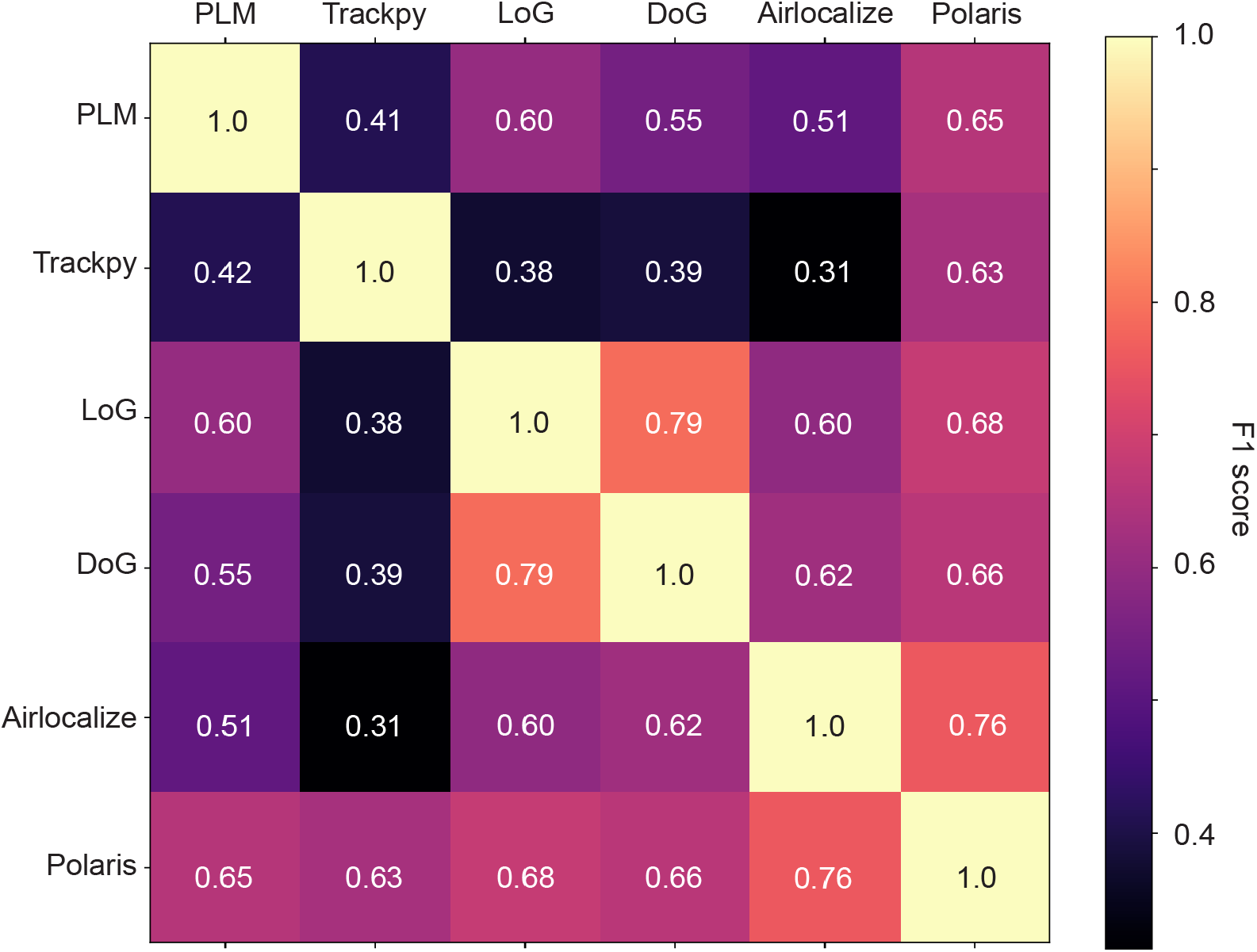
Quantification of agreement between Polaris’ deep learning model and different classical spot detection methods. The benchmarked methods include maximum-intensity filtering (PLM), the Crocker-Grier centroid-finding algorithm (Trackpy), Laplacian of Gaussian (LoG), difference of Gaussians (DoG), Airlocalize, and Polaris.

### 6.6 Supplementary Note 6: Benchmarking the receptive field of Polaris’ spot detection model

**Figure S5:**
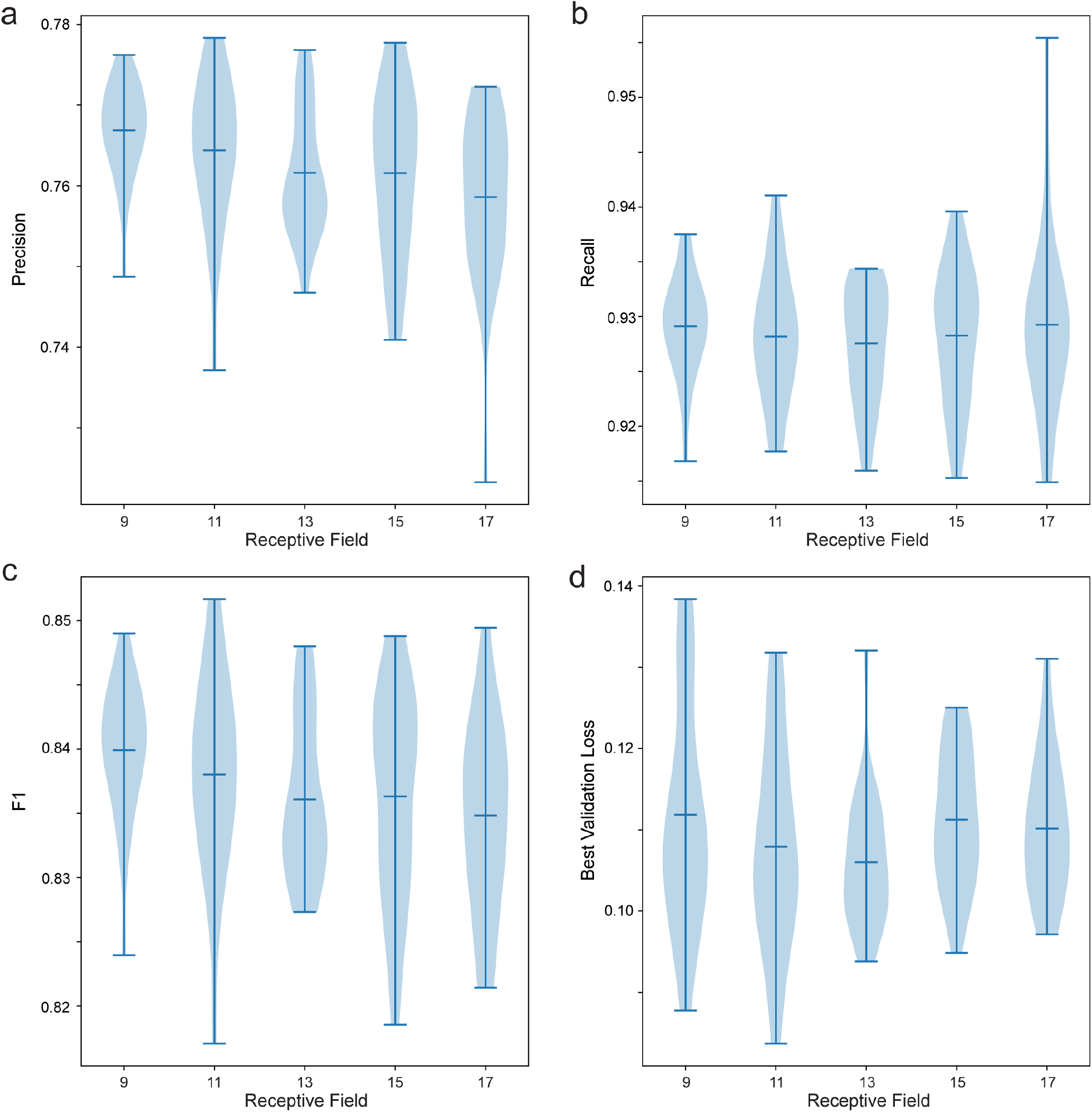
Benchmarking the receptive field parameter of Polaris’ spot detection model. (a-d) Violin plot quantifying the performance metrics ((a) precision, (b) recall, (c) F1, (d) best validation loss during training) for models trained with different values for receptive field of Polaris’ spot detection model. n=24 trained models per receptive field condition.

### 6.7 Supplementary Note 7: Benchmarking Polaris’ spot detection model on simulated images with ranging spot characteristics

**Figure S6:**
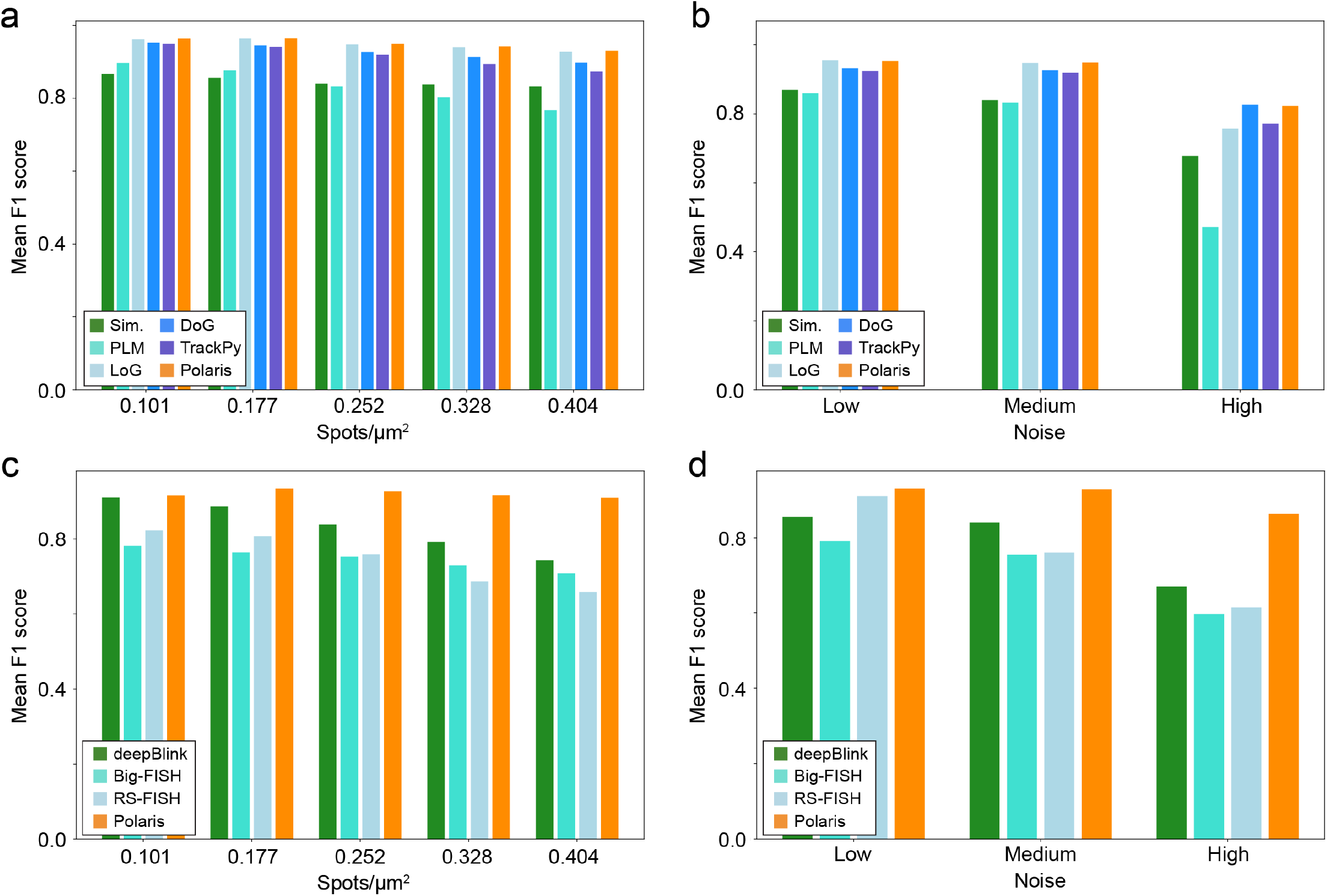
Benchmarking model performance on simulated spot images with a range of spot intensities and densities. (a) Performance quantification for models with Polaris’ deep learning architecture trained with various datasets predicting on images with a range of spot density. (b) Performance quantification for models with Polaris’ deep learning architecture trained with various datasets predicting on images with a range of levels of simulated noise. The low noise condition corresponds to a signal-to-noise ratio of greater than approximately 16. The medium noise condition corresponds to a SNR of approximately 8-15. The high noise condition corresponds to a SNR of approximately 3-7. (c) Performance quantification for various spot detection methods on images with a range of spot densities. (d) Performance quantification for various spot detection methods on images with a range of levels of simulated noise. The SNR ranges for each noise condition are the same as those in (b).

### 6.8 Supplementary Note 8: Robustness of Polaris’ barcode decoding method to dropout

**Figure S7:**
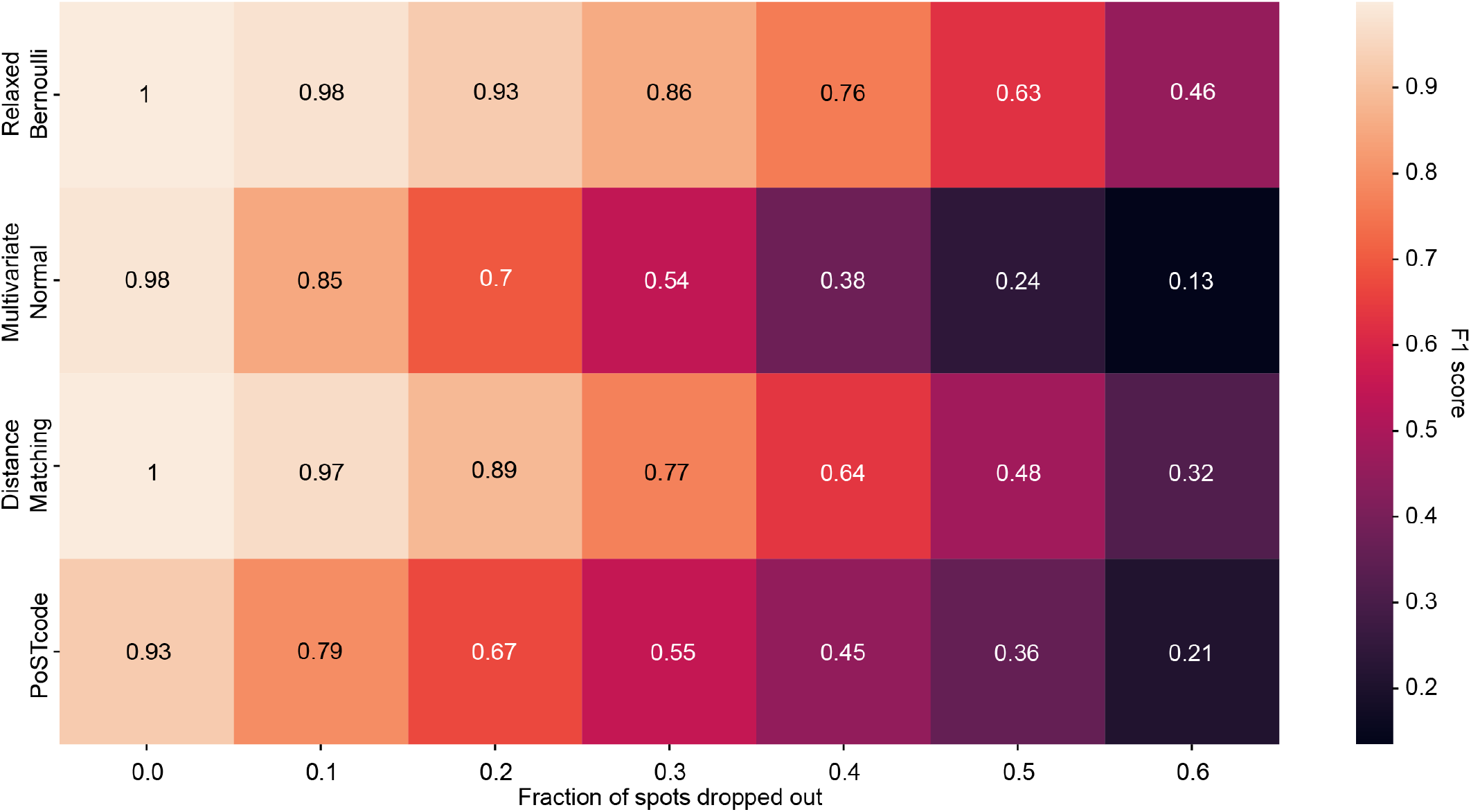
Benchmarking of the robustness of gene decoding methods to dropout. Quantification of F1 score for four barcode decoding methods (a graphical model of relaxed Bernoulli distributions, a graphical model of multivariate normal distributions, Hamming distance matching, and PoSTcode) for simulated barcode pixel values with a range of dropout rates.

### 6.9 Supplementary Note 9: Benchmarking Polaris’ performance on various multiplexed FISH image sets

**Figure S8:**
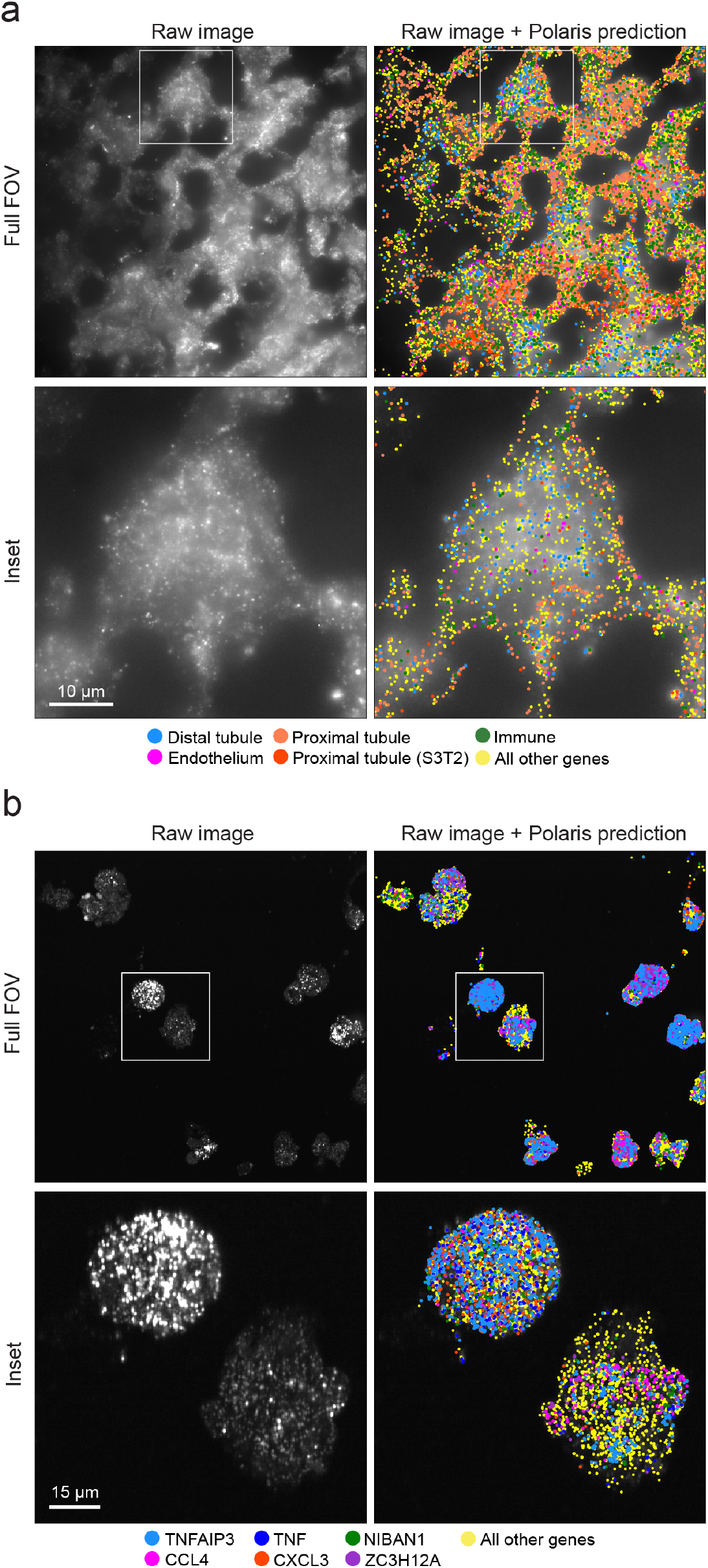
Demonstration of Polaris’ performance on a MERFISH and seqFISH data. (a) Example Polaris prediction for a MERFISH experiment in a mouse kidney tissue sample (Liu, et al. 2022). (b) Example Polaris prediction for a seqFISH experiment in a macrophage cell culture sample. The spot colors of the Polaris prediction in (a,b) denote the predicted gene identities. The inset image location is defined by the white box in the full field of view (FOV).

**Figure S9:**
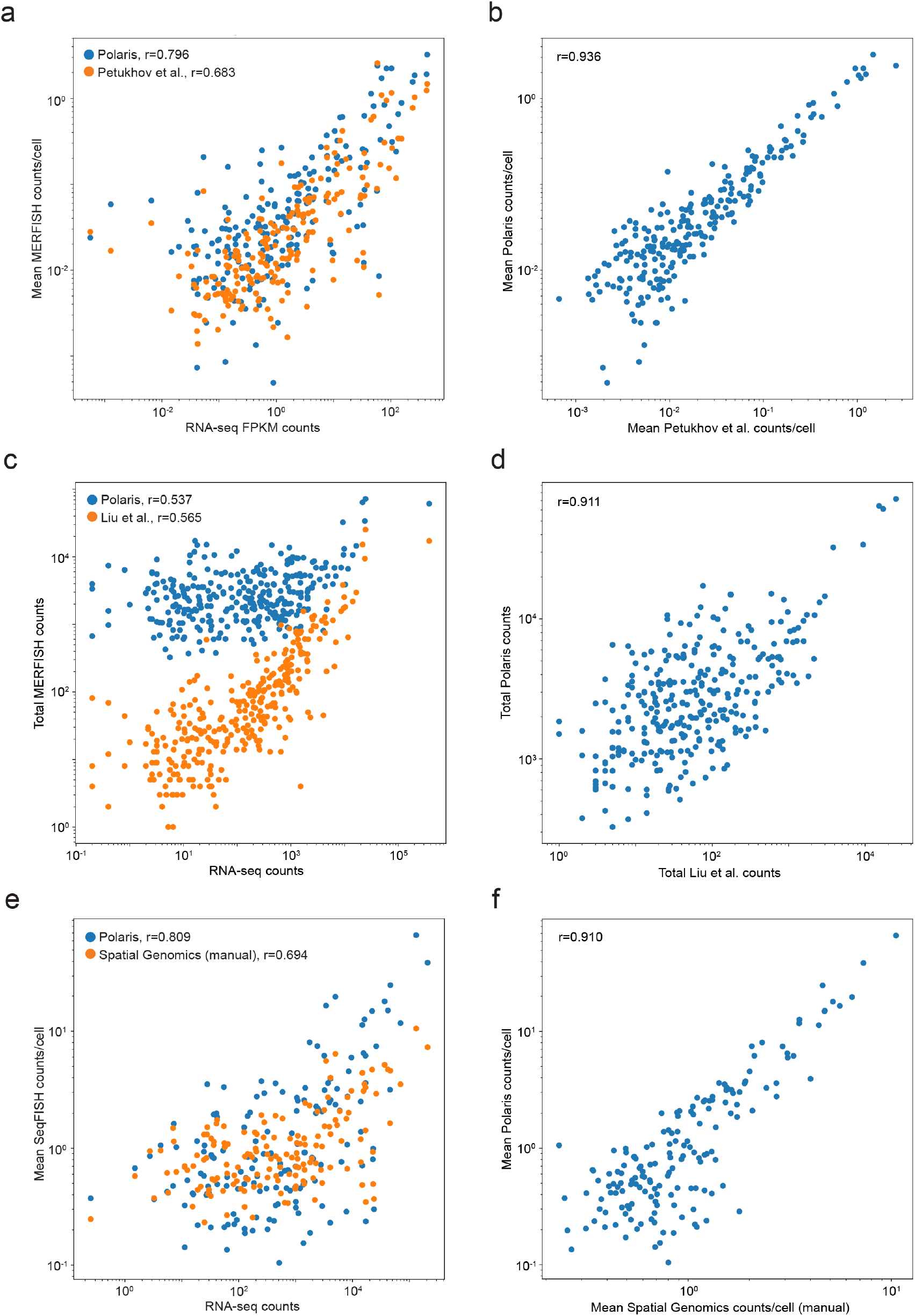
Correlation of Polaris’ quantification of MERFISH data with other quantification methods. (a,c) Scatter plot plotted in logspace, comparing gene expression counts quantified with MERFISH with counts measured with RNA-seq. (b,d) Scatter plot plotted in logspace, comparing previously published MERFISH gene expression counts with counts quantified with Polaris. (e) Scatter plot plotted in logspace, comparing mean gene expression counts per cell quantified with seqFISH with counts measured with RNA-seq. (f) Scatter plot plotted in logspace, comparing mean gene counts per cell obtained by manual analysis of seqFISH data with gene counts quantified with Polaris.

### 6.10 Supplementary Note 10: Demonstration of Polaris’ performance decoding an ISS barcode library

**Figure S10:**
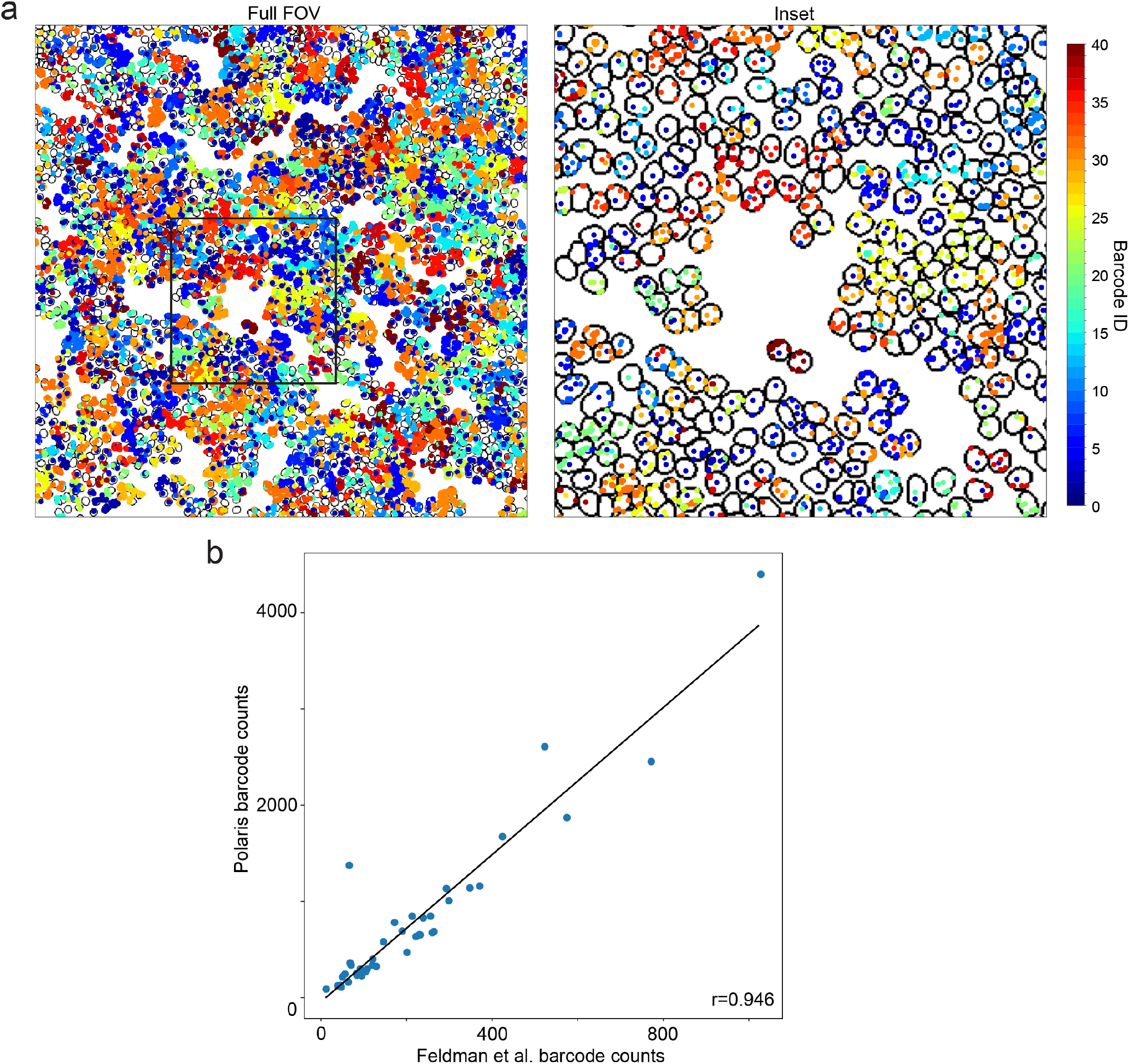
Demonstration of Polaris’ performance on an ISS dataset in HeLa cells. (a) Example Polaris prediction for the ISS sample. The spot colors correspond with barcode identities. The inset location is defined by the black box in the full field of view (FOV). (b) Scatter plot correlating total counts for each barcode decoded by the original published analysis with counts quantified by Polaris (r=0.946).

